# Preferential activation of proprioceptive and cutaneous sensory fibers compared to motor fibers during cervical transcutaneous spinal cord stimulation: A computational study

**DOI:** 10.1101/2022.02.02.478757

**Authors:** Roberto M. de Freitas, Marco Capogrosso, Taishin Nomura, Matija Milosevic

**Affiliations:** Graduate School of Engineering Science, Department of Mechanical Science and Bioengineering, Osaka University, Japan; Department of Neurological Surgery, University of Pittsburgh, Pittsburgh, USA; Rehab and Neural Engineering Labs, University of Pittsburgh, Pittsburgh, USA; Department of Bioengineering, University of Pittsburgh, Pittsburgh, USA

**Keywords:** cervical, transcutaneous spinal cord stimulation, upper-limb, computational simulation, finite element method

## Abstract

**Objective:** Cervical transcutaneous spinal cord stimulation (tSCS) is a promising technology that can support motor function recovery of upper-limbs after spinal cord injury. Its efficacy may depend on the ability to recruit sensory afferents and convey excitatory inputs onto motoneurons. Therefore, understanding its physiological mechanisms is critical to accelerate its development towards clinical applications. In this study, we used an anatomically realistic computational model of the cervical spine to compare α-motor, Aα-sensory, and Aβ-sensory fiber activation thresholds and activation sites.

**Approach:** We developed a tridimensional geometry of the cervical body and tSCS electrodes with a cathode centred at the C7 spinous process and an anode placed over the anterior neck. The geometrical model was used to estimate the electric potential distributions along motor and sensory fiber trajectories at the C7 spinal level using a finite element method. We implemented dedicated motor and sensory fiber models to simulate the α-motor and Aα-sensory fibers using 12, 16, and 20 μm diameter fibers, and Aβ-sensory fibers using 6, 9, and 12 μm diameter fibers. We estimated nerve fiber activation thresholds and sites for a 2 ms monophasic stimulating pulse and compared them across the fiber groups.

**Main results:** Our results showed lower activation thresholds of Aα- and Aβ-sensory fibers compared with α-motor fibers, suggesting preferential sensory fiber activation. We also found no differences between activation thresholds of Aα-sensory and large Aβ-sensory fibers, implying they were co-activated. The activation sites were located at the dorsal and ventral root levels.

**Significance:** Using a realistic computational model, we demonstrated preferential activation of dorsal root Aα- and Aβ-sensory fibers compared with ventral root α-motor fibers during cervical tSCS. These findings suggest high proprioceptive and cutaneous contributions to neural activations during cervical tSCS, which inform the underlying mechanisms of upper-limb functional motor recovery.

## 1. Introduction

Cervical transcutaneous spinal cord stimulation (tSCS) has recently emerged as a non-invasive rehabilitation technology for recovery of upper-limb motor function after spinal cord injury [1–3]. Neuromodulation of cervical spinal neural circuitries may occur when cervical tSCS is combined with supraspinal descending drive, promoting neuroplasticity across the damaged spinal regions [1,4,5]. Specifically, it has previously been suggested that these rehabilitation effects may depend on the activation of sensory fibers during tSCS [6–9]. On the other hand, direct activation of motor fibers may be adverse to the rehabilitation efficacy as it could lead to multiple-muscle contractions [10–12]. However, little is known about which specific groups of fibers are activated during cervical tSCS. Better understanding of the neural activation targets could therefore provide implications for rehabilitation and inform therapeutic functional parameter selection for cervical tSCS.

The lack of knowledge on what groups of motor and sensory fibers are co-activated during cervical tSCS also limits our interpretation and analysis of the spinally motor evoked potentials in surface electromyography (EMG) [13–16]. For instance, in previous cervical tSCS experimental studies, the activation of Ia-sensory fibers was inferred by post-activation depression of the motor evoked potentials during application of paired stimuli [13–15,17]. Specifically, post-activation depression was conditioned by two stimulating pulses with equal amplitudes that were sequentially delivered with short (e.g., 50 ms) interstimulus intervals [13–16]. A decrease in the amplitude of the second motor evoked potential in relation to the first has been suggested to be primarily caused by a suppression of monosynaptic connectivity between Ia-sensory fibers and α-motoneurons after the first stimulus [13–16]. Additionally, when the second motor evoked motor potentials were not fully suppressed, this suggested possible activations of fibers groups other than the Ia-sensory fibers [13,14]. These other fiber groups could include large diameter fibers, such as α-motor and Ib-sensory fibers, as previously suggested in experimental studies [13,14,18,19]. Moreover, using experimental recordings and computational simulations, it was also suggested that cutaneous Aβ-sensory fibers could be co-activated in non-human primates during epidural cervical spinal cord stimulation [10]. Taking into consideration the practical feasibility of invasively recording neural activities in humans, the activation of sensory and motor fibers during cervical tSCS at a neural level has not been confirmed.

Computational simulations can play an important role in better understanding the physiological mechanisms underlying motor and sensory fiber activations during spinal cord stimulation at a neural level [10,20–23]. Previous studies have coupled simulations with nerve fiber models to reproduce realistic membrane dynamics in response to extraneural electrical fields, thereby accurately predicting activations of motor and sensory fibers [10,20–22,24]. For instance, accurate representations of the proprieties of membrane fibers (e.g., membrane resistance and capacitance) according to the fiber diameters may help reproduce preferential activation of large diameter nerve fibers over the smaller ones during electrical stimulation [10,23,25]. In previous computational studies examining lumbar tSCS, activations of motor fibers in the ventral roots and sensory fibers in the dorsal roots were analyzed during monophasic stimulation pulse [20,21]. Cervical tSCS computational models, on the other hand, have not yet been developed, primarily owning to the later development of this technology in the rehabilitation of upper-limbs.

The anatomy of the cervical body and the lower trunk spinal cord around the lumbar vertebrae is considerably different [26–28], implying that nerve fiber activation mechanisms during lumbar and cervical tSCS are also different. For instance, the ventral and dorsal rootlets are shorter and more obtuse at the cervical body compared with the lower trunk [26,27]. Moreover, the curvature of the spinal cord, as well as the shape of the vertebral bones also differ between the two anatomical regions [29]. These geometrical differences influence not only in the curvature of the trajectories of motor and sensory nerve fibers, but also the current flow across the spinal canal where the nerve fibers cross [20,30]. Indeed, the substantial anatomical differences between lumbar and cervical body may critically affect outcomes of cervical tSCS compared to lumbosacral stimulation [31]. Therefore, the results obtained with simulations of lumbar tSCS may not be directly translatable to cervical tSCS, and specific computational models are required to better understand the neural activation mechanisms underlying cervical stimulation.

Considering the lack of direct evidence from previous experimental studies in analyzing nerve fiber activation during cervical tSCS, we used computational simulations to better understand the activations of motor and sensory fibers at the neural level. In this study, a cervical tSCS model was developed to simulate activations of α-motor and Aα-sensory fibers using 12, 16, and 20 μm diameters [21,32], as well as Aβ-sensory fibers using 6, 9, and 12 μm diameters [10]. As such, the recruitment of motor, proprioceptive (Ia- and Ib-sensory fibers), and cutaneous fibers were estimated by simulating, the α-motor, Aα-sensory, and Aβ-sensory fibers, respectively [33]. Specifically, the cathode electrode was configured at the C7 spinal process while the anode electrode was configured on the anterior neck, following electrode configurations of cervical tSCS experimental studies [13–16]. The minimum current intensities injected in the anode electrode that are necessary to activate the nerve fibers (activation thresholds) and the location where the action potentials were initiated (activation sites) were compared across different diameter size motor and sensory fibers.

Taking into account the evidence shown by previous experimental studies, which suggested that spinally motor evoked potentials during cervical tSCS are largely elicited by the transsynaptic activation of sensory fibers [13–15], we hypothesized that Aα-sensory fibers would be preferentially activated compared with α-motor fibers. In other words, we expect that the Aα-sensory fibers would have lower activation thresholds compared with the α-motor fibers due to their different physiological properties [32,34–36]. Moreover, considering the inverse recruitment order of nerve fibers observed experimentally during electrical stimulation [25], we also hypothesized that the Aβ-sensory fibers would be least excitable due their smaller range of diameters. In other words, we expected that Aβ-sensory fibers would have the highest activation thresholds compared with α-motor and Aα-sensory fibers. This is contrary to what was previously shown by epidural spinal stimulation modeling studies [10]. However, the larger dimensions of the cervical tSCS electrodes, as well as their distance from the targeted fibers compared with epidurally placed electrodes are likely to cause different activation mechanism between the non-invasive and invasive spinal stimulation approaches [20].

## 2. Methods

We first developed a tridimensional geometry of the cervical body and stimulation electrodes based on MRI reconstructed volume of the human body and anatomical values from the literature (see section 2.1). Using this geometry, we then estimated the electric potential distribution along the trajectories of motor and sensory fibers of the C7 spinal level using the FEM (see section 2.2). Computational axon fiber models were used to simulate the activation of α-motor, Aα-sensory, and Aβ-sensory fibers when extraneural electric potentials were applied (see section 2.3). Finally, we computed the activation thresholds, which are the minimum stimulation intensities necessary to activate the nerve fibers, as well as the locations where the action potentials were initiated, to compare the motor and sensory fibers of different diameters (see section 2.4).

### 2.1. Development of an anatomically realistic cervical tSCS geometrical model

We developed a tridimensional model representing anatomical structures of the cervical body and stimulation electrodes using Autodesk Inventor Professional 2021 (Autodesk Inc., USA) as illustrated in Figure 1. The geometry of the cervical body was designed based on the MRI reconstructed volume of the “virtual family” from a 34 year old male model “Duke” [28], and anatomical values from the literature [13,14,26,27,37–42]. Specifically, the geometry model included 27 components: white matter and gray matter [26]; dorsal and ventral rootlets, and the respective roots, from the right and left sides at C7 spinal level [27,37]; dorsal root ganglions [40]; left and right C7 spinal nerves designed until the clavicle level [39]; cerebrospinal fluid [38]; C5, C6, C7, and T1 spinal bones with C5-C6, C6-C7, and C7-T1 inter-vertebral spinal disks [28]; epidural fat [28]; general cervical body [28]; back muscles [28]; fat [41]; skin [42]; a cathode electrode (5×5 cm) placed over C7 spinal process [13–15]; and an anode electrode (7.5×10 cm) placed on the anterior side of the neck [13–16].

**Figure 1:**
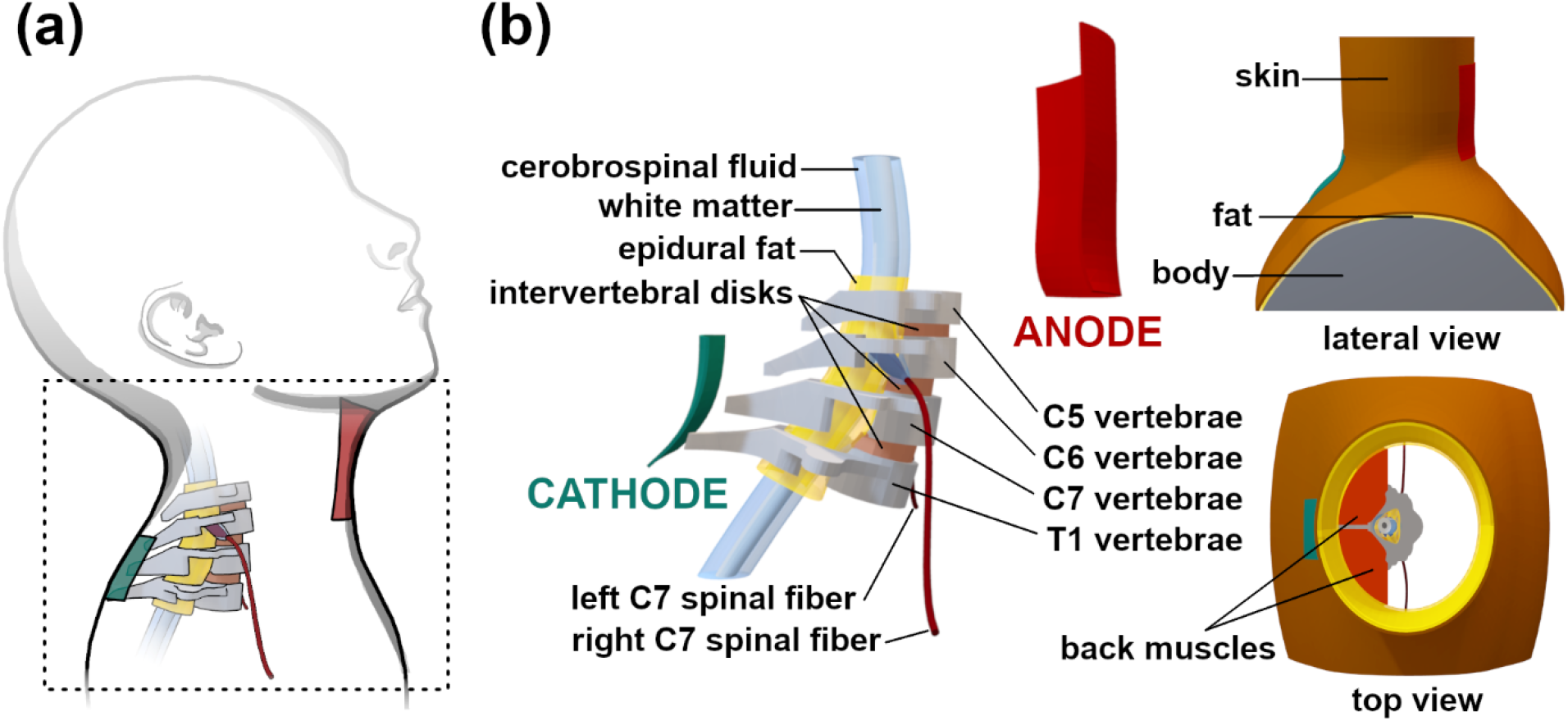
(a) Illustration of the cervical body configuration during cervical transcutaneous spinal cord stimulation (tSCS) considered for the development of the tridimensional model. A cathode (green) and an anode (red) electrodes are arranged in a configuration used by previous cervical tSCS experimental studies [13–15]. Specifically, the cathode electrode configured over the C7 spinal process and the anode electrode configured over the anterior neck. (b) Tridimensional model of the cervical body developed based on MRI-derived realistic anatomical proportions. On the left, most of the model components surrounding the spinal canal are shown. The gray matter and dorsal and ventral roots are not visible. In the lateral view, the fat, body and skin structures are indicated. In the top view, the back muscles enclosed between the fat and vertebra structures are also shown, while the body structure is not visible.

### 2.2. Electric potential distribution estimation

The tridimensional model was meshed in COMSOL Multiphysics (v.5.6, COMSOL Inc., USA) with approximately 8.5 million tetrahedral elements. We assigned each component of the geometry an electrical conductivity *σ* as indicated in Table 1, consistent with previous studies [20–22]. Anisotropic electrical conductivities (longitudinal x transversal) of the white matter and back muscles were implemented using the Curvilinear Coordinates toolbox in COMSOL [10]. Specifically, the diffusion method was used, and their bottom and top surfaces were defined with the inlet and outlet conditions, respectively [10]. All components were considered as purely resistive, and a quasi-static assumption was adopted to calculate the distribution of electric potentials *V*(***x***) at any point ***x*** of the domain during stimulation [20,21,43,44]. Insulating boundary condition was applied to all external surfaces of the geometry [22]. Dirichlet boundary condition was applied to cathode electrode with *V*(***x***) = 0. Neuman boundary condition was applied to the anode electrode such that a 1 A electric current was injected [10,22]. The Laplace equation ∇. (*σ*(***x***)∇*V*(***x***)) = 0, with the electrical conductivity *σ*(***x***) not constant for the white matter and muscle, was solved with the FEM in COMSOL to obtain the electric potential distributions across the tridimensional geometry.

**Table 1:** Conductivity values, in siemens per meter (S/m), of the materials assigned to the components of the geometrical model of the cervical body (gray matter, white matter, ventral/dorsal roots and rootlets, dorsal root ganglia, C7 spinal nerves, cerebrospinal fluid, epidural fat, C5-C6, C6-C7 and C7-T1 intervertebral disks, C5, C6, C7 and T1 vertebra, back muscles, general cervical body, skin and fat) and the electrodes (i.e., anode and cathode electrodes). For white matter and muscles, anisotropic conductivities were assigned to longitudinal (L) and transversal (T) components.

| *Material* | *Conductivity $\sigma$ (S/m)* |
| --- | --- |
| *Gray matter* | 0.276 |
| *White matter* | L: 0.6; T: 0.083 |
| *Spinal nerve, root, dorsal root ganglion and rootlet* | 0.1432 |
| *Cerebrospinal fluid* | 1.7 |
| *Epidural fat* | 0.04 |
| *Intervertebral disk* | 0.6 |
| *Spinal bone* | 0.02 |
| *Muscle* | L: 0.5; T: 0.08 |
| *Body* | 0.25 |
| *Skin* | 0.0025 |
| *Fat* | 0.04 |
| *Electrode* | 0.01 |

Legend: L – longitudinal; T – transversal.

### 2.3. Axon models

A sensory and a motor axon fiber models developed by Gaines and colleagues [32] were used to simulate the dynamics of the membrane potentials during cervical tSCS [22,45]. The Gaines models [32] are derived from the model developed by McIntyre, Richardson, and Grill (MRG) [46], and implements adjustments to the specific channel gating parameters to account for motor and sensory fibers physiological differences [34–36]. For both motor and sensory fiber models, the nerve fiber consists of nodal and internodal segments, as illustrated in Figure 2a. Each node of Ranvier is represented by a single segment (NODE), and each internode by ten segments: two paranodal myelin attachment segments (MYSA); two paranodal main segments (FLUT); and six internodal segments (STIN) [32,46]. The Gaines models [32] include fast K^+^ channels in the node segments, and fast K^+^, slow K^+^, leak and hyperpolarization activated cyclic-nucleotide gated (HCN) channels in the internodal segments, which accounts for a more realistic representation of ion channels of motor and sensory fibers [47]. The membrane and myelin sheath conductance and capacitance values are defined according to the diameter and length dimensions of the nodal and internodal segments (NODE, MYSA, FLUT, and STIN). In turn, the NODE diameter, STIN diameter, STIN length, length between two NODE, number of myelin lamellae and STIN length, were linearly extrapolated from [46] for each axon fiber diameter size that was simulated (i.e., for 6, 9, 12, 16, and 20 μm), consistent to pervious studies [21]. The Gaines motor and sensory axon fiber models and the MGR model were implemented in Python 3.9 using NEURON 8.0 [48].

**Figure 2:**
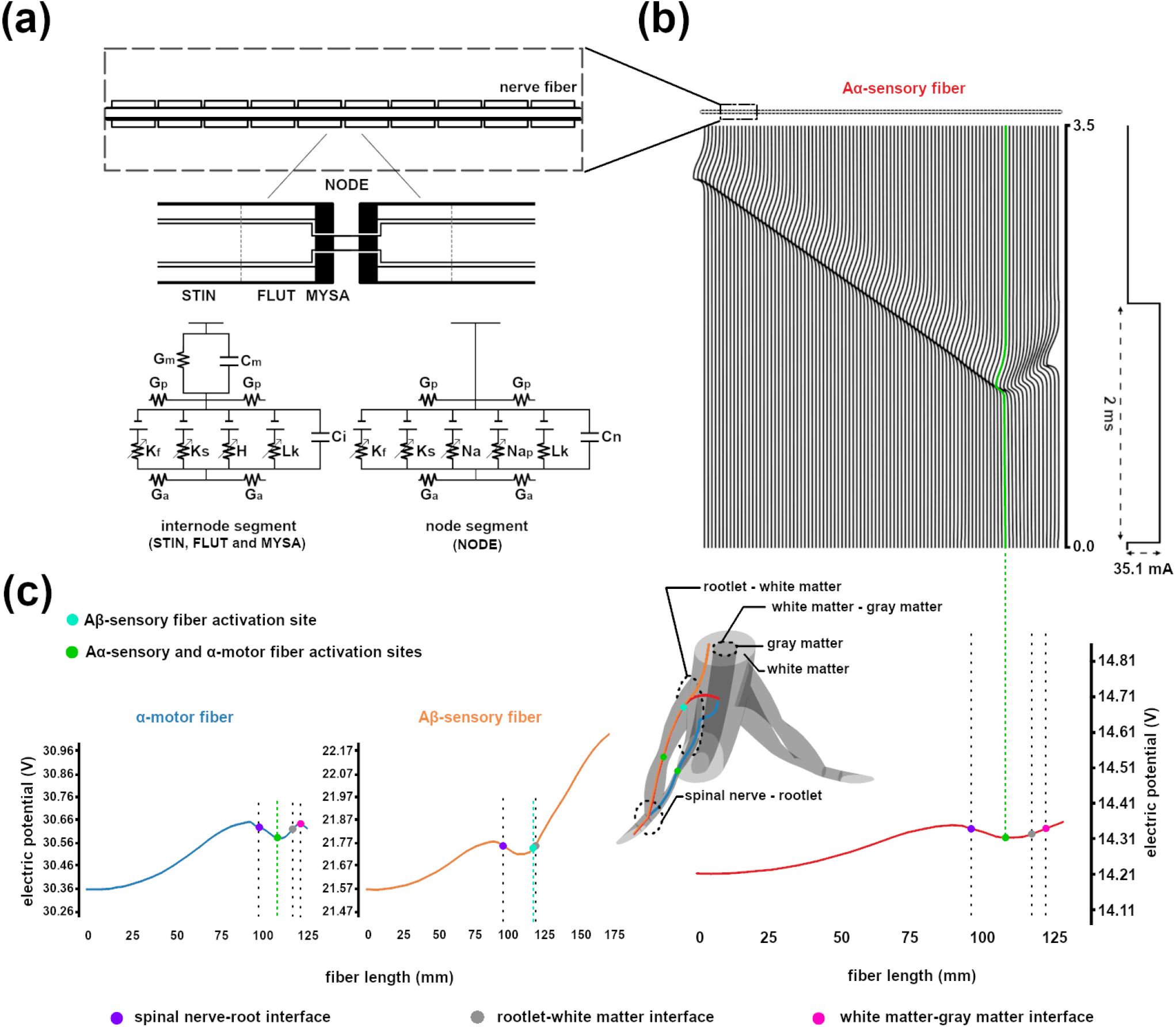
(a) Compartment model of myelinated nerve fiber membrane proposed by Gaines et al. [32] to simulate motor and sensory fibers. Each node (NODE) is represented by one segment, and each internode by ten segments: two MYSA, two FLUT and six STIN. The node and internode segments are composed of fast K^+^ (K_f_), slow K^+^ (K_s_), fast Na^+^ (Na), persistent Na^+^ (Na_p_), leak current (L_k_) and HCN (H) channels. Nodal (C_n_), internodal (C_i_) and myelin (C_m_) capacitances, as well as axoplasmic (G_a_), periaxonal (G_p_) and myelin (G_m_) conductances are also represented in this model. For details, see [32,46]. (b) At the top, a representative response of the membrane potentials in all node segments simulated with Gaines model. The representative response is from a 16 μm diameter Aα-sensory fiber with the trajectory entering the gray matter. The minimum stimulation intensity necessary to activate the fiber was 38.1 mA for a 2 ms pulse width. The location where the action potential was initiated, referred to as activation site, was located at the middle portion of the dorsal root, indicated by the green trace. (c) On the right side, the electric potential distribution along the Aα-sensory fiber is shown with its activation site indicated by a green circle. On the left side, representative electric potential distributions along a 16 μm diameter α-motor fiber (blue) and a 9 μm diameter Aβ-sensory fiber (orange) are shown with its corresponding activation sites indicated by green and cyan circles, respectively. The stimulation amplitude defined as minimum stimulation intensity necessary to activate the α-motor and Aβ-sensory fibers were 75.0 mA and 53.3 mA, respectively, for a 2 ms pulse width. The location at interface between the spinal nerve and the rootlets, at the interface between the rootlets and the white matter, and at the interface between the white and gray matters are indicated by purple, gray and pink circles, respectively.

### 2.4. Simulation conditions

#### Gaines model simulations

The distribution of electric potentials was estimated along the α-motor, Aα-sensory, and Aβ-sensory fiber trajectories, which are shown in Figure 2. Specifically, different fiber trajectories were defined along the left and right C7 spinal nerves from the clavicles level until inside the spinal cord. At each dorsal (sensory fibers) and ventral (motor fibers) roots, three different trajectories were defined spanning across the middle portion of the rootlet structures. At the interface between the ventral rootlets and the white matter, each motor fiber was further defined in three other pathways leading towards the center of the gray matter. Moreover, at the interface between the dorsal rootlets and the white matter, each Aα-sensory fiber trajectory was defined in three other pathways towards the center of the gray matter, while each Aβ-sensory fiber trajectory was defined in three ascending pathways in the dorsal column [49], consistent with a previous study [10]. In total, 18 trajectories were defined for each of the α-motor, Aα-sensory, and Aβ-sensory fibers, including the right and left sides (i.e., 3 segments at the roots level x 3 segments entering the white matter x 2 sides = 18 trajectories).

To estimate the activation thresholds, which are the minimum stimulation intensity required to activate the fibers, we scaled the distributions of electric potentials along the motor and sensory fiber trajectories and applied them as extraneural potentials to the axon fiber models [50,51]. Specifically, for each fiber, the extraneural potential distribution was computed as the minimum level required for fiber activation using a bisection algorithm [21,22]. The extraneural potential distributions were applied by simulating a rectangular monophasic stimulation pulse with a 2 ms pulse duration, consistent with previous cervical tSCS experimental studies [13–15]. It is noteworthy that the stimulation current applied at the anode electrode and the electrical potential in the domain, *V*(***x***), are linearly related. Therefore, the scaling factor estimated to elicit fiber activation corresponds to a fraction of the 1 A initially simulated at the anode electrode with FEM simulations. Furthermore, we also estimated the locations along the trajectories where the action potentials were initiated at the activation threshold intensity, i.e., activation sites.

The α-motor, Aα-sensory, and Aβ-sensory fibers were simulated using the motor and sensory fiber models proposed by Gaines et al. [32]. The α-motor and Aα-sensory fibers with trajectories entering the gray matter were simulated using 12, 16, and 20 μm diameters [33], which were used in previous simulation studies [21,32]. Specifically, this range of fiber diameters covers the sizes of Ia- and Ib-sensory fibers, as well as large diameter α-motor fibers [33]. The Aβ-sensory fibers ascending the dorsal column were simulated using 6, 9, and 12 μm diameters, consistent with previous simulation studies [10]. These diameter sizes are within the range corresponding to the group of Aβ-sensory fibers and also ascending branches of Ia-sensory fibers, which were considered to be two-thirds of the diameters of the dorsal root segments [49]. The 18 trajectories that were simulated for each fiber type caused the electric potential distribution along each fiber to be slightly different, thereby introducing variations in the activation thresholds, which allowed for statistical comparison.

#### MRG model stimulation

To examine the methodological considerations used by Danner and colleagues [21], we also used the MRG model [46] to estimate the activation thresholds of the motor and sensory fibers, which were differentiated only by their diameter sizes. In agreement with their methodology, only motor and sensory fibers entering the gray matter were simulated (i.e. the same 18 trajectories defined for α-motor and Aα-sensory fibers), while the trajectories ascending to the dorsal column from the dorsal roots were omitted [21]. Specifically, the sensory fibers were simulated with the 16 μm diameters and motor fibers with the 14 μm diameters [21].

### 2.6. Statistics

Since normality of the data could not confirmed for all analysis groups using the Shapiro-Wilk test, nonparametric tests were used. First, the Kruskal-Wallis test was used to compare activation thresholds of α-motor, Aα-sensory and Aβ-sensory fibers that were pooled across their respective fiber diameters (*fiber type*: α-motor, Aα-sensory, and Aβ-sensory). Moreover, the Kruskal-Wallis test was also used to compare the activation thresholds across groups of motor and sensory fibers with different diameters to examine possible co-activations (*fiber groups*: α-motor_12_, α-motor_16_, α-motor_20_, Aα-sensory_12_, Aα-sensory_16_, Aα-sensory_20_, Aβ-sensory_6_, Aβ-sensory_9_, and Aβ-sensory_12_). When significant results were found for the Kruskal-Wallis test, multiple-pairwise comparisons were performed with Bonferroni corrections. Additionally, the Mann-Whitney test was used to compare the activation thresholds of motor and sensory fibers estimated using the MRG model (*MRG fibers*: MRG_sensory16_ and MRG_motor14_). Statistical tests were performed using SPSS (IBM, Armonk, NY) and the significance level was set to α=0.05.

## 3. Results

### 3.1. Gaines model motor and sensory fibers activation thresholds and sites

We used the motor and sensory fiber models developed by Gaines and colleagues [32] to simulate α-motor, Aα-sensory, and Aβ-sensory fibers. The α-motor and Aα-sensory were simulated with 12, 16, and 20 μm diameters, whereas Aβ-sensory fibers were simulated with 6, 9, and 12 μm diameters. For each fiber group, the activation thresholds and sites were estimated using 18 trajectories (see sections 2.4 and 3.2), shown in Figure 3. We identified the activation sites at the activation threshold intensity level for each simulated fiber (Figure 3). The Kruskal-Wallis test was used to compare between α-motor, Aα-sensory, and Aβ-sensory fibers (*fiber type*: α-motor, Aα-sensory, and Aβ-sensory). Moreover, the Kruskal-Wallis test was used to compare the activation thresholds of motor and sensory fibers with different diameter sizes (*fiber groups*: α-motor_12_, α-motor_16_, α-motor_20_, Aα-sensory_12_, Aα-sensory_16_, Aα-sensory_20_, Aβ-sensory_6_, Aβ-sensory_9_, and Aβ-sensory_12_).

**Figure 3:**
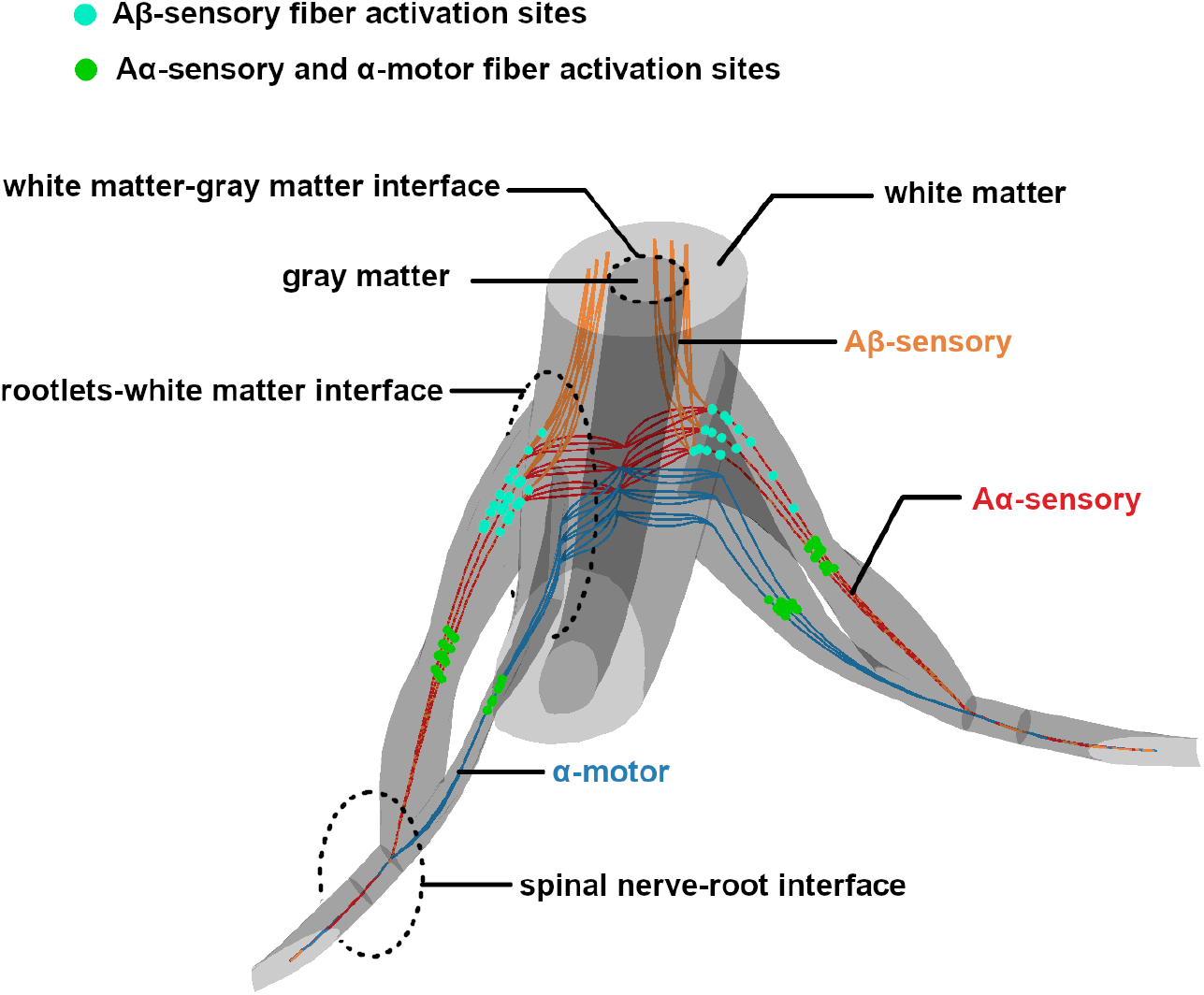
Activation sites of all Aα-sensory (red traces) and α-motor fibers (blue traces) are indicated by green circles at the middle portion of the ventral and dorsal roots. The trajectories of these fibers are defined to cross through the white matter and enter the gray matter. The activation sites of Aβ-sensory fibers (orange traces) are indicated by cyan circles around the interface between the dorsal rootlets and the white matter. The trajectories of Aβ-sensory fibers are defined to ascend the dorsal column when they enter the white matter. In total, 18 α-motor, 18 Aα-sensory, and 18 Aβ-sensory fibers were simulated with the myelinated nerve fiber model developed by Gaines et al. [32].

#### Activation sites

A representative simulated response of the membrane potentials in all node segments during cervical tSCS is presented in Figure 2b. In this illustrative example, we show the activation of a Aα-sensory fiber. Specifically, the 16 μm diameter Aα-sensory fiber that was simulated using the Gaines model for a 2 ms stimulation pulse with amplitude activation threshold amplitude (see section 2.4.1.). An action potential was initiated approximately at the middle portion of the dorsal root, corresponding to a valley point in the extraneural electric potential distribution applied along the fiber length, as illustrated on the right side of Figure 2c. Analogous to Aα-sensory fibers, the activation sites of α-motor fibers were located in the middle portion of the ventral root, as exemplified on the left side of Figure 2c. Contrary to the α-motor and Aα-sensory fibers, the activation sites of Aβ-sensory fibers were located closer to the interface between the dorsal roots and white matter, as illustrated on the left side of Figure 2c. Consistent with these representative outcomes, the activation sites of all simulated motor (α-motor) and sensory (Aα- and Aβ-sensory) fibers in our study are shown in Figure 3. Indeed, the activation sites for α-motor and Aα-sensory always corresponded to locations at the level of the roots for both the left and right sides. In contrast, the activation sites for Aβ-sensory fibers were consistently located closer to the interface between rootlets and white matter, where these fibers curved towards ascending the pathway in the dorsal column.

#### Activation thresholds

Comparison results of the activation thresholds across different fiber diameters sizes (*fiber type*: α-motor, Aα-sensory, and Aβ-sensory) are summarized in Figure 4a. Our results showed significant main effects between the *fiber type* factors (χ^2^(2,162)=70.90, *p*<.001; α-motor: 86.11±22.41 mA, Aα-sensory: 39.72±12.58 mA, and Aβ-sensory: 72.91±45.23 mA). Multiple pairwise comparisons with Bonferroni corrections showed that the activation thresholds of Aα-sensory fibers were significantly lower compared to α-motor (*p*<.001) and Aβ-sensory fibers (*p*<.001), as well as that activation thresholds of Aβ-sensory fibers were significantly lower compared to α-motor fibers (*p*=.001) (Figure 4a).

**Figure 4:**
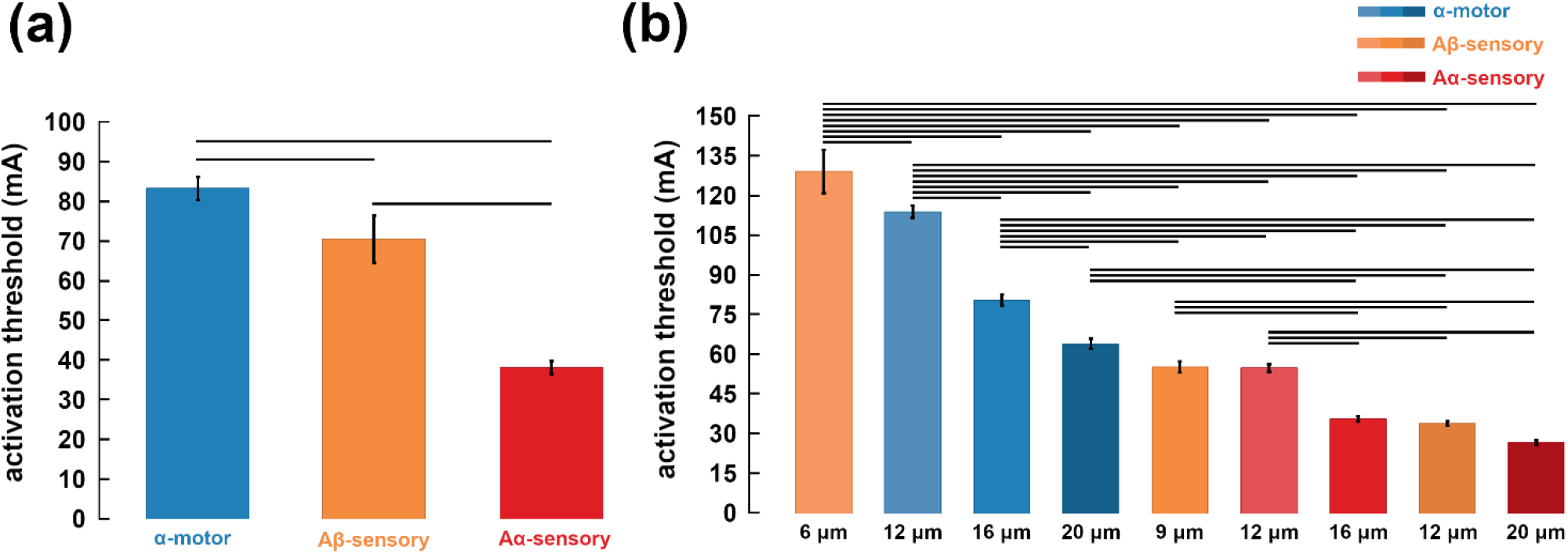
(a) The minimum cervical transcutaneous spinal cord stimulation intensity, referred to activation threshold (mA), necessary to activate α-motor (blue), Aα-sensory (red), and Aβ-sensory (orange) fibers was compared using the Kruskal-Wallis test. The α-motor and Aα-sensory fibers were both simulated with 12, 16 and 20 μm diameters, whereas Aβ-sensory fibers were simulated with 6, 9 and 12 μm diameters. (b) The activation thresholds of the fibers were grouped by diameter sizes and compared using the Kruskal-Wallis test. In total, 9 groups of fibers (α-motor, Aα- and Aβ-sensory fibers simulated with 3 diameters sizes) were compared, each with 18 different trajectories. All the fibers were simulated with the myelinated nerve fiber membrane proposed by Gaines et al. [32]. Statistically significant differences found between the groups are indicated by the horizontal bars. The bar plot bins indicate the means of the activation thresholds of each analysis group, and the black vertical bars their standard errors.

Comparison of co-activation of fibers with different diameter sizes across groups of motor and sensory fibers (*fiber groups*: α-motor_12_, α-motor_16_, α-motor_20_, Aα-sensory_12_, Aα-sensory_16_, Aα-sensory_20_, Aβ-sensory_6_, Aβ-sensory_9_, and Aβ-sensory_12_) are summarized in Figure 4b. Our results showed significant main effects between *fiber groups* factors (χ^2^(8,162)=150.76, *p*<.001; α-motor_12_: 113.47±9.78 mA, αmotor_16_: 80.52±8.83 mA, α-motor_20_: 64.34±8.01 mA, Aα-sensory_12_: 55.25±6.16 mA, Aα-sensory_16_: 36.34±4.53 mA, Aα-sensory_20_: 27.58±3.36 mA, Aβ-sensory_6_: 128.47±33.93 mA, Aβ-sensory_9_: 55.72±8.57 mA, and Aβ-sensory_12_: 34.54±3.67 mA). Multiple pairwise comparisons with Bonferroni corrections showed that the activation thresholds between a majority of the fibers were significantly different (*p*<.05). Notably, the Aβ-sensory_12_, Aα-sensory_16_, and Aα-sensory_20_ fibers, as well as α-motor_20_, α-motor_16_, Aβ-sensory_9_, and Aα-sensory_12_ fibers were not significantly different (*p*>.05) (Figure 4b).

### 3.3. MRG model simulations of motor and sensory fibers

We used the MRG nerve fiber model [46] and the considerations used by Danner and colleagues [21] to simulate the motor and sensory fibers with 14 and 16 μm diameters [21], respectively. The activation thresholds were estimated using the same 18 trajectories as the α-motor and Aα-sensory fibers (see sections 2.4 and 3.2), which are shown as blue and red in Figure 3. The Mann-Whitney test was used to compare the motor and sensory activation thresholds simulated with the MGR model (*MRG fibers*: MRG_sensory16_ and MRG_motor14_).

Comparison between MRG_sensory16_ (16 μm diameter) and MRG_motor14_ fibers (14 μm diameter) are summarized in Figure 5. Our result showed that activation thresholds of MRG_sensory16_ and MRG_motor14_ fibers were not significantly different (*U*=105.50, *p*=.074; MRG_motor14_: 34.15±4.25 mA and MRG_sensory16_: 31.41±4.00 mA).

**Figure 5:**
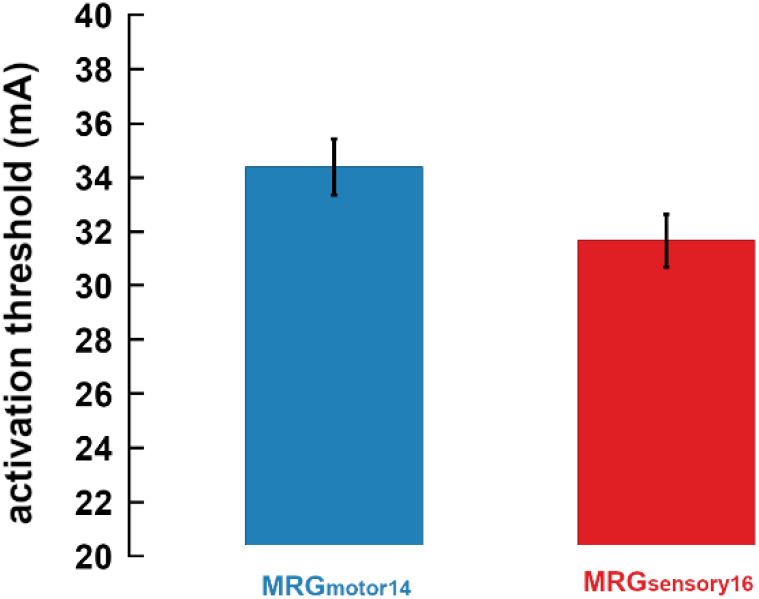
The minimum cervical transcutaneous spinal cord stimulation intensity, referred as activation threshold (mA), necessary to activate motor (blue) and sensory fibers (red) were compared using the Mann-Whitney test. Motor fibers were simulated with 14 μm and sensory fibers with 16 μm using the myelinated nerve fiber membrane proposed by McIntyre et al., [46]. No statistically significant differences were found. The bar plot bins indicate the means of the activation thresholds of each analysis group, and the black vertical bars their standard errors.

## 4. Discussions

We developed a computational model to better understand the neural activation of motor and sensory fibers during cervical tSCS. We found that Aα-sensory and large Aβ-sensory fibers (9 and 12 μm diameters) had lower activation thresholds compared with α-motor fibers (Figure 4b). Moreover, the activation sites of α-motor and Aα-sensory fibers were located approximately at the middle portion of the ventral and dorsal roots, respectively, whereas the activation sites of Aβ-sensory were located around the interface between the dorsal rootlets and the white matter. Despite anatomical differences between the cervical and lumbar body, our findings also showed preferential sensory fiber activations at the cervical level as previously demonstrated for lumbar tSCS [21]. Notably, our findings also suggest a large contribution of cutaneous (i.e., Aβ-sensory) fiber activations during cervical tSCS, despite their relatively small sizes. Taken together, our study elucidates the neural activation mechanisms and provide implications for the therapeutic application of cervical tSCS.

### 4.1. Motor and sensory fiber activations

Previous experimental studies in humans have used post-activation depression of distally recorded EMG responses to investigate weather the recruitment of Ia-sensory fibers affected the cervical tSCS evoked motor potentials [13–17]. In agreement with our hypothesis that Aα-sensory fiber group (representing Ia- and Ib-sensory fibers) would have lower activation thresholds compared with α-motor fibers, our findings confirmed preferential activation of Aα-sensory compared with α-motor fibers (Figure 4a). These results are consistent with the mechanism inferred from post-activation depression, which suggested strong contribution of Ia-sensory fiber activation to spinally motor evoked potentials [13,14]. Despite not including specific considerations for distinguishing Ia- and Ib-fibers in our simulations, they are likely to be co-activated due to their overlapping range of diameters [33]. While the activation of Ib-sensory fibers may activate polysynaptic pathways that inhibit the of spinally motor evoked potentials, they are unlikely to have large a effect on post-activation depression [17]. Furthermore, we showed that large Aβ-sensory fibers ascending from the dorsal roots toward the dorsal column (9 and 12μm diameters) were activated with lower stimulation intensities compared with the α-motor fibers (Figure 4b). Comparison across all diameters seems to indicate that Aβ fibers are activated with higher stimulation intensities compared with Aα fibers (Figure 4a). However, a closer inspection of activations thresholds between different diameter sizes revealed that Aα- and Aβ-sensory fibers were co-activated across most fiber sizes (Figure 4b). Contrary to what was suggested in previous experimental studies, our simulations therefore showed an sizable contribution of cutaneous fibers to the motor evoked potentials during cervical tSCS [17,52]. While cutaneous afferents may contribute to motor evoked potentials, they may also convey divergent excitatory potentials to relevant spinal nodes and significantly contribute to the rehabilitative potential of non-invasive spinal stimulation [53–55]. Taken together, our results showed preferential recruitment of Aα- and Aβ-sensory fibers compared with α-motor fibers, suggesting large contribution of proprioceptive and cutaneous activations during cervical tSCS.

It is well known that nerve fibers with larger diameters are preferentially activated compared with those with smaller diameters when extraneural electrical potentials are applied [25,56]. However, contrary to our hypothesis that Aβ-sensory fibers would have the highest activation thresholds due to their smaller size, we showed that large Aβ fibers (9 and 12 μm diameters) were recruited at similar stimulation intensities as the Aα fibers (12, 16, and 20 μm diameters) (Figure 4b). These results are consistent with the findings of Greiner and colleagues [10] who showed co-activation of Aα and Aβ fibers using computer simulations in non-human primate models [10]. Despite the smaller range of diameter sizes and that cervical tSCS electrodes are located farther away from the targeted sensory dorsal roots, our results nonetheless also suggest a sizable contribution of cutaneous Aβ-sensory fibers during non-invasive cervical spinal stimulation. The co-activation Aα and Aβ fibers could be explained by their electric potential distributions, as illustrated in Figure 2c. After the interface between the dorsal roots and the white matter, the electrical potential distribution of Aβ-sensory fibers is increased as the fibers project further away from the cathode and approach the anode electrode. The Aα-sensory fiber electrical potential distributions do not have such large changes after entering the white matter. For instance, Aα-sensory fibers simulated using 12 μm diameter seemed to have lower activation thresholds compared with the Aβ-sensory fibers which were simulated using the same diameter size (Figure 4b). These fibers were different only in their trajectories after entering the spinal cord, highlighting the large influence of trajectories on defining the electric potential distributions and consequent activation thresholds. Therefore, the pathways defining the projection of different fibers seem to compensate for the different diameters of the Aα fibers and Aβ fibers. Overall, our simulations showed co-activation of cutaneous and proprioceptive nerve fibers despite their different diameters, which may also possibly suggest that non-invasive cervical tSCS can activate some common fiber types as cervical epidural spinal stimulation [10].

### 4.2. Activation sites during cervical tSCS

Differences between the cervical and lower trunk body anatomy, such as the geometrical shape of the vertebra as well as the curvature of dorsal and ventral rootlets at the level where they enter the vertebra, imply that results obtained from lumbar tSCS simulation studies [20,21] may not directly translate to cervical tSCS. These studies showed that activation sites of motor and sensory fibers were located approximately at the point where they enter the white matter and exit the spinal canal [20]. In agreement with lumbar tSCS results, our simulations showed activation sites of Aβ-sensory fibers located around the interface between the dorsal rootlets and the white matter [20,21]. However, activation sites of α-motor and Aα-sensory fibers were located approximately at the middle portion of the rootlets structure (Figure 3). It is noteworthy that α-motor and Aα-sensory fibers were simulated using different membrane ion channel properties (see section 2.4.1) but were defined along similar trajectories that terminate inside the gray matter (Figure 3). As discussed already, the differences between the Aβ-sensory activation sites and that of α-motor and Aα-sensory fibers can likely to be attributed to their different trajectories, and the consequent electric potential distribution along the respective fibers (Figure 2c) [57]. The anatomical structures used in our simulations, which consequently affected the geometry of motor and sensory nerve fiber trajectories, could explain activation site differences between cervical and lumbar tSCS simulations obtained previously [20,21].

### 4.3. Comparisons with previous simulations and experimental data

In the current study, the Gaines models were used for simulating motor and sensory fibers, accounting for their physiological differences [34–36]. Contrary to the results obtained by Danner and colleagues [21], the activation thresholds of motor (14 μm diameter) and sensory (16 μm diameter) fibers simulated with the MRG model [46] were not different in our current study, despite the activation threshold for the sensory fibers appearing to be lower (Figure 5). It is not clear why these specific diameter sizes were used by Danner et al. [21]. However, it should be noted that increasing the difference between the motor and sensory diameters within a physiological range, e.g., using 13 μm diameter for motor and 17 μm diameter for sensory, can yield larger differences between their activation thresholds. We used the motor and sensory fibers proposed by Gaines et al. [32], which allowed us to directly compare motor and sensory fibers with the same diameters (Figure 4b). Notably, our findings also expanded the work of Danner and colleagues [21] by considering trajectories of Aβ-sensory fibers with pathways defined from the dorsal roots towards acceding levels of the dorsal column.

The activation intensities estimated using the Gaines models in our simulations are numerically compatible with previous experimental studies [14,15]. For instance, de Freitas et al. [14] showed that motor thresholds of upper-limb proximal muscles were in the range between 35 and 70 mA, which is compatible with the range of the activation thresholds of Aα- and Aβ-sensory fibers (i.e., activation thresholds between 30 and 60 mA in Figure 4b). Similarly, Sasaki et al. [15] showed that the stimulation intensity range for eliciting post-activation depression were approximately 57.0 ± 4.0 mA, also in the range of the thresholds obtained in our simulations (Figure 4b). Therefore, our simulation results obtained herein have a close correspondence with previous experimental studies, supporting the robustness of our model.

### 4.4. Implications for therapeutic application of cervical tSCS

Our results demonstrated the contribution of proprioceptive fibers during cervical tSCS, which has been suggested to contribute to neuromodulation [4,6,9] and motor rehabilitation of spinal cord injured patients [1,3,5]. Specifically, proprioceptive fiber activation compensates for the scarce signal transmissions during motor tasks in spinal cord injured patients [6]. During attempted voluntary movements, Ia-sensory fiber activation could convey excitatory inputs onto motoneurons to amplify the motor outputs, which could result in neuroplasticity across the spinal lesioned areas [6,9]. Our finding also showed preferential activation of cutaneous fibers, indicating the potential use of cervical tSCS for the treatment of other clinical conditions. Considering the importance of cutaneous activation to the sensorimotor rehabilitation of stroke survivors [53,54], it is possible that cervical tSCS may also be effective in post-stroke rehabilitation. Moreover, activation of Aβ-sensory fibers could activate inhibitory circuits at the spinal and supraspinal levels which attenuate nociceptive signals [58,59], implying possible benefits for chronic pain treatment.

This reasoning is supported by the recent evidence showing utility of transcutaneous spinal direct current stimulation in chronic pain care [55]. Irrespective of the intended application, the stimulation intensity of cervical tSCS should be configured just around the level to transsynaptically elicit motor potentials through selective activation of proprioceptive and cutaneous fibers, i.e., without direct motor fiber activation [3,4,12,60]. In clinical settings, this may be achieved by the stimulation intensity around the level for eliciting paresthesias [60], which could be attained by Aβ-sensory fibers activation [10,61]. However, it is important to note that other parameter selection such as the duration of stimulation [15] and voluntary involvement during stimulation [4] are likely required for effective neuromodulation during cervical spinal stimulation. Taken together, our findings imply that the activation of Aα- and Aβ-sensory fibers during cervical tSCS likely plays a role in motor rehabilitation of spinal cord injured patients [1,3,5], also suggesting potential utility of this technology for treatment of other clinical conditions such as sensorimotor impairment caused by stroke [53,54] and for chronic pain [55].

### 4.5. Limitations and future directions

A limitation of the our tridimentional geometry is related to some simplifications adopted by the model. For instance, the back muscles were designed to fill the distance between the vertebra and fat, whereas the gray matter was apprximated using a clyndrical shape (Figure 3). On the other hand, the rootlets and the spinal canal curvature, which are the main neural tragets in our study, were developed with detailed considerations such that the dimensional curvature of the nerve fibers were fully representative (Figure 3). Considering the distance between the stimulating electrodes and the fibers where the electric potentials were calculated, these geometrical abstractions are considered acceptable, or perhaps even more detailed than previous simulations [20,21]. Future studies are nontheless warranted to expand the complexity of the current cervical tSCS model in order to study the activation of nerve fibers at different spinal levels and using different electrode configurations.

## 5. Conclusions

We developed a computational model of cervical tSCS to analyze neural activation of α-motor, Aα-, and Aβ-sensory fibers. Our results showed that dorsal root proprioceptive Aα and cutaneous Aβ fibers were co-activated at lower stimulating intensities compared with the ventral root α-motor fibers. Preferential activation of sensory fibers can be attributed to their physiological proprieties and different trajectories which define the electric potential distribution along their extent. Notably, our study demonstrated sizable cutaneous contributions during non-invasive spinal stimulation, along with co-activated with the proprioceptive fibers. Understanding these neural activation mechanisms is critical for defining parameters to selectively activate dorsal root sensory fibers, an important consideration for rehabilitation effectiveness of spinal stimulation.

## Acknowledgments

R.M.F. is supported by the Engineering Science for the 21^st^ Century scholarship granted by the Japanese Ministry of Education, Culture, Sports, Science and Technology (MEXT). This project was funded by the Japan Society for the Promotion of Science Grants-in-Aid for Scientific Research - KAKENHI (Grant numbers: 20K19412).

